## Supplementary material for "Anthracycline-induced cardiotoxicity associates with a shared gene expression response signature to TOP2-inhibiting breast cancer drugs in cardiomyocytes": S1 Appendix

### Table of contents

#### Figures

S1 Fig: Cardiomyocytes can be generated at high purity across six individuals.

S2 Fig: Dose-response curves are reproducible across replicate cardiomyocyte differentiations from the same individual.

S3 Fig: Cancer drugs that decrease cardiomyocyte viability induce cellular stress.

S4 Fig: Treatment with ACs at a dose of 0.5 μM for 48 hours induces effects on cardiomyocyte viability.

S5 Fig: RNA-seq sample quality is equivalent across individuals, treatments, and time points.

S6 Fig: RNA-seq samples cluster by treatment type, timepoint, and individual.

S7 Fig: PC1 associates with drug treatment and treatment time, while PC2 associates with individual.

S8 Fig: Thousands of gene expression changes are induced in response to TOP2i treatment over 24 hours.

S9 Fig: ACs affect expression of nearly half of all expressed genes after 24 hours of treatment.

S10 Fig: A small number of genes respond to a single drug only.

S11 Fig: Stringently-identified drug-specific response genes are enriched in biological processes.

S12 Fig: Most genes that are differentially expressed in response to treatment at three hours are also differentially expressed at 24 hours.

S13 Fig: Four gene expression signatures capture the response to TOP2i over time.

S14 Fig: AC treatments induce a small number of gene expression variation changes across individuals.

S15 Fig: TOP2i treatments induce expression changes in some genes that correlate with cardiotoxicity.

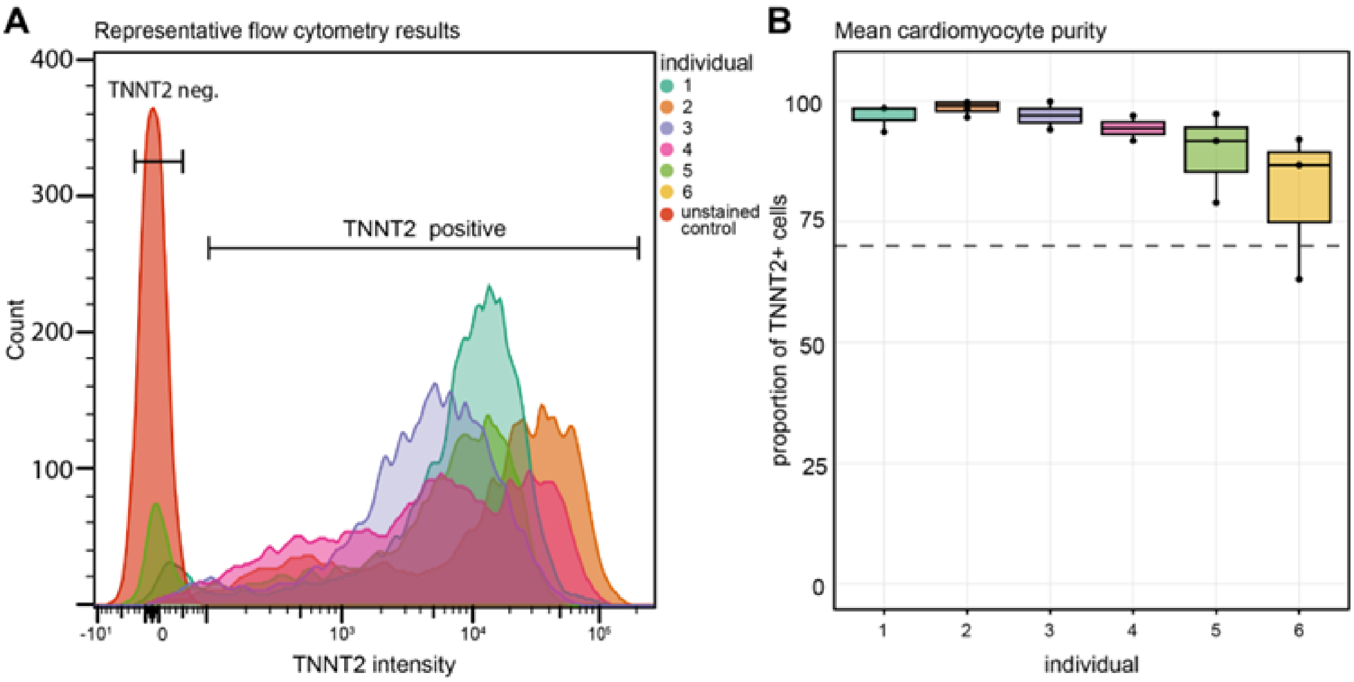


**Figure S1: Cardiomyocytes can be generated at high purity across six individuals. (A)** Representative image of flow cytometry data indicating the proportion of TNNT2 positive cells in one differentiation experiment for each individual based on the fluorescent intensity of the phycoerythrin-labeled TNNT2 antibody and a sample of unlabeled iPSC-CMs (red cell population). **(B)** Percentage of cells that are positive for expression of TNNT2 for each individual. Data representative of three independent differentiation experiments used for the two drug dose-response curves, and RNA and cell culture media collection. The dashed line represents high-purity iPSC-CMs (> 70% TNNT2 positive).

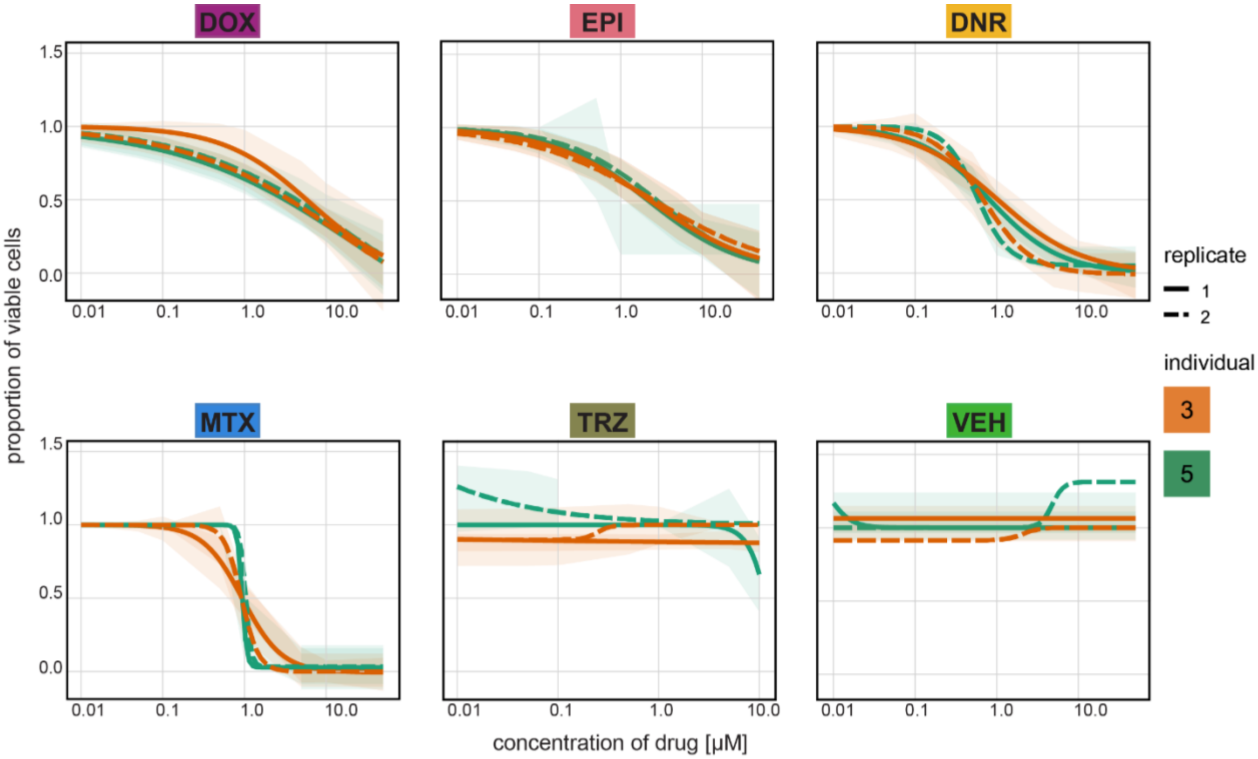


**Figure S2: Dose-response curves are reproducible across replicate cardiomyocyte differentiations from the same individual.** Proportion of viable cardiomyocytes following exposure to increasing concentrations of each drug. Cell viability in replicate one (solid line) and two (dashed line) in Individual three (orange), and Individual five (green) was assessed following 48 hours of drug treatment. Viability was determined at each drug concentration in quadruplicate, and the mean value was selected for generation of the dose-response curves using a four-point log-logistic regression with the upper asymptote set to 1. Shading represents the 95% confidence interval from the regression analysis for Individual three (light orange) and Individual five (light green).

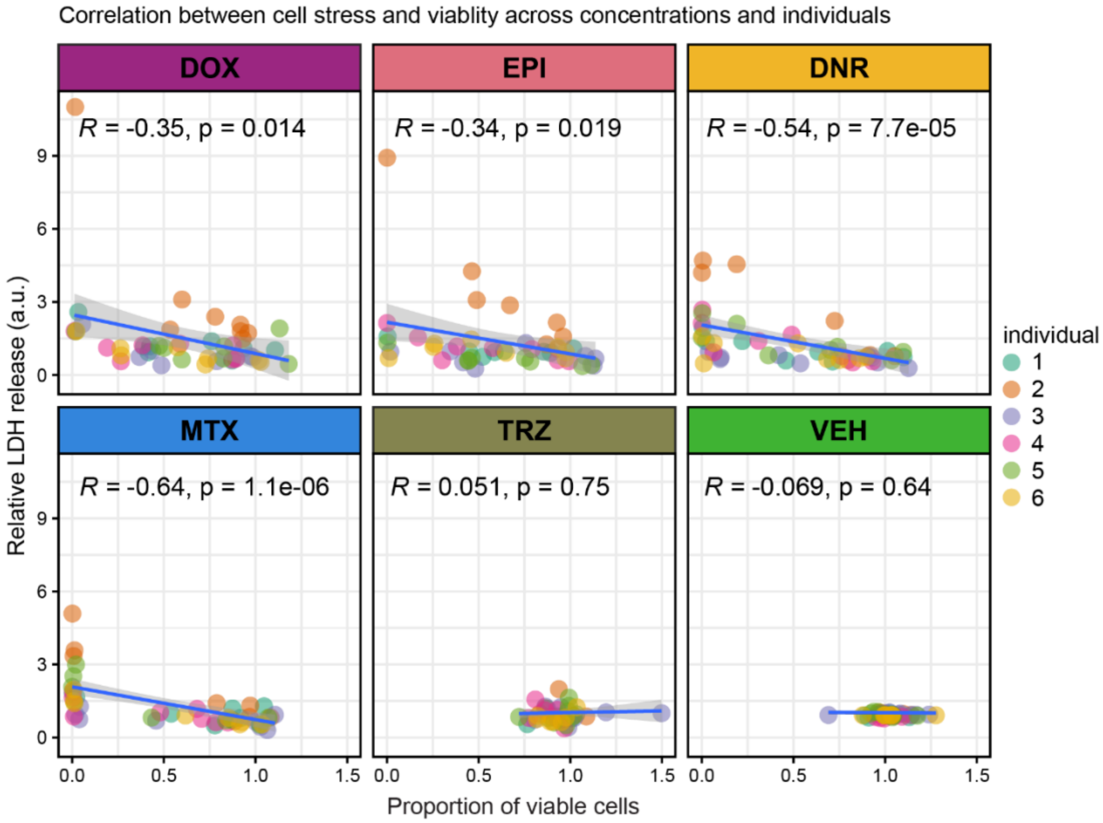


**Figure S3: Cancer drugs that decrease cardiomyocyte viability induce cellular stress.** Pearson correlation between cardiomyocyte viability following drug treatment at eight different concentrations for 48 hours, and the level of lactate dehydrogenase released into the cell culture media across individuals. Data points are colored by individual (1,2,3,4,5,6).

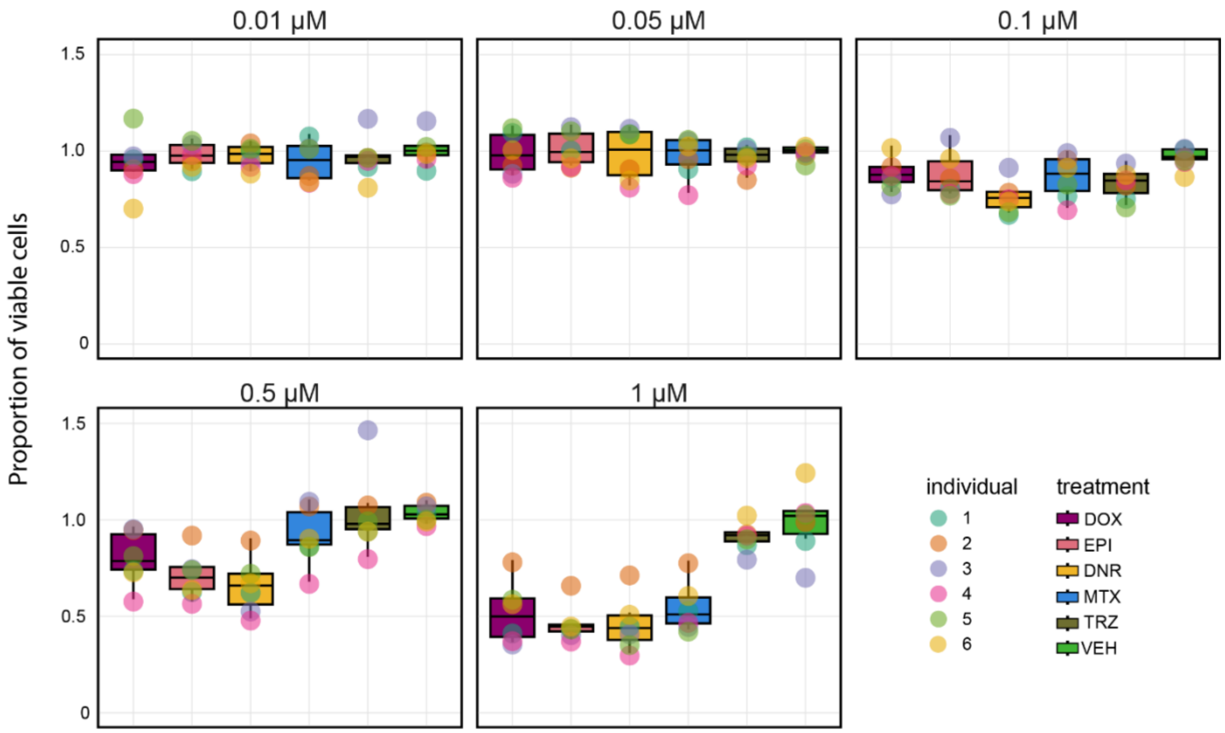


**Figure S4: Treatment with ACs at a dose of 0.5 μM for 48 hours induces effects on cardiomyocyte viability.** Proportion of viable cells following treatment with each drug (DOX: mauve; EPI: pink; DNR: yellow; MTX: blue; TRZ: dark green; VEH: light green) in each individual (1,2,3,4,5,6) at five sub-micromolar drug concentrations.

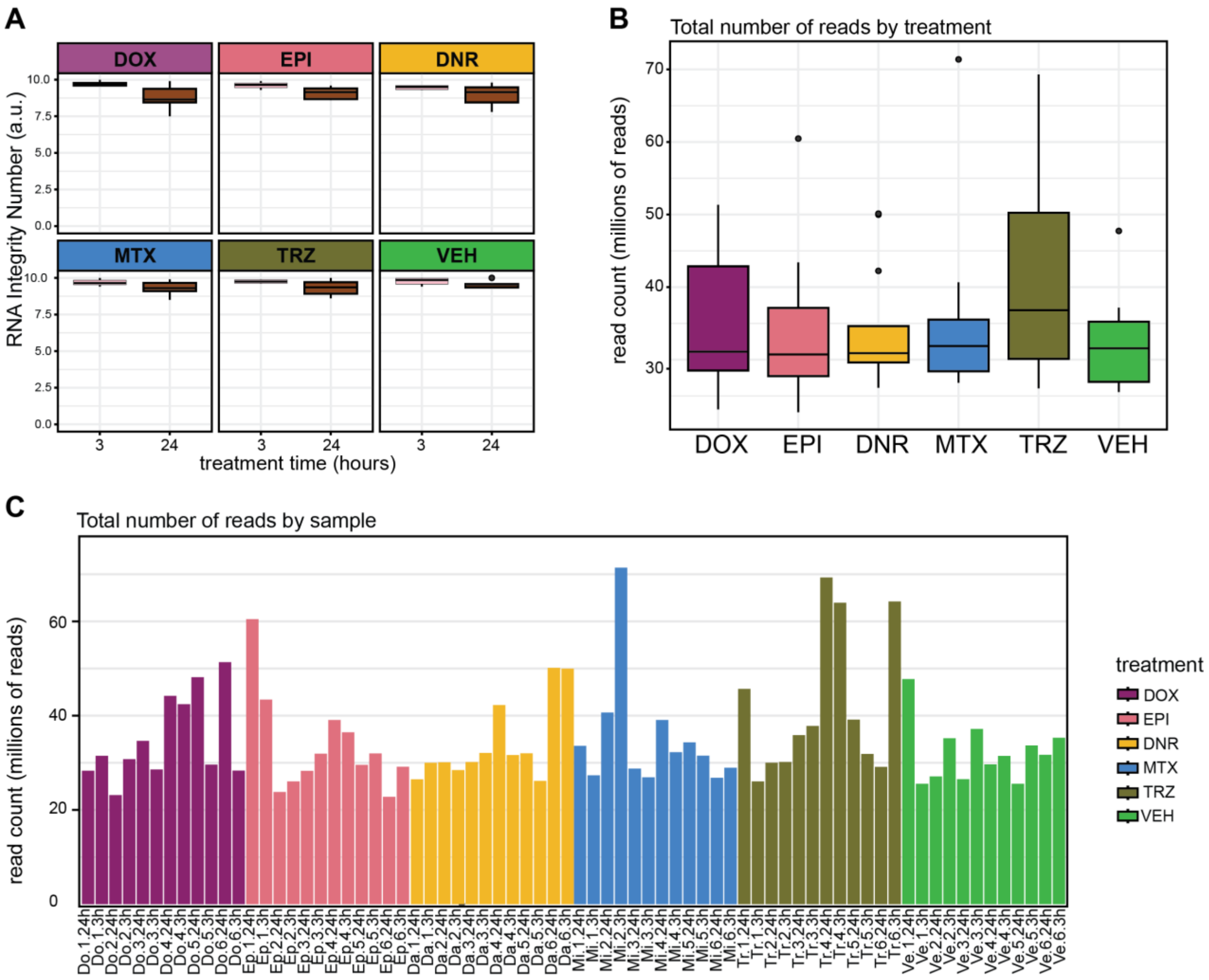


**Figure S5: RNA-seq sample quality is equivalent across individuals, treatments, and time points. (A)** RNA integrity score for each sample categorized by drug type and drug treatment time. Data inclusive of six individuals. **(B)** Total number of RNA-sequencing reads categorized by treatment type (DOX: mauve; EPI: pink; DNR: yellow; MTX: blue; TRZ: dark green; VEH: light green). Each drug treatment category includes data from six individuals across two time points. **(C)** Total number of RNA-seq reads for each of the 72 samples. Each sample is denoted by drug.individual.timepoint, eg. DOX-treatment in Individual one collected at 24 hours (Do.1.24h).

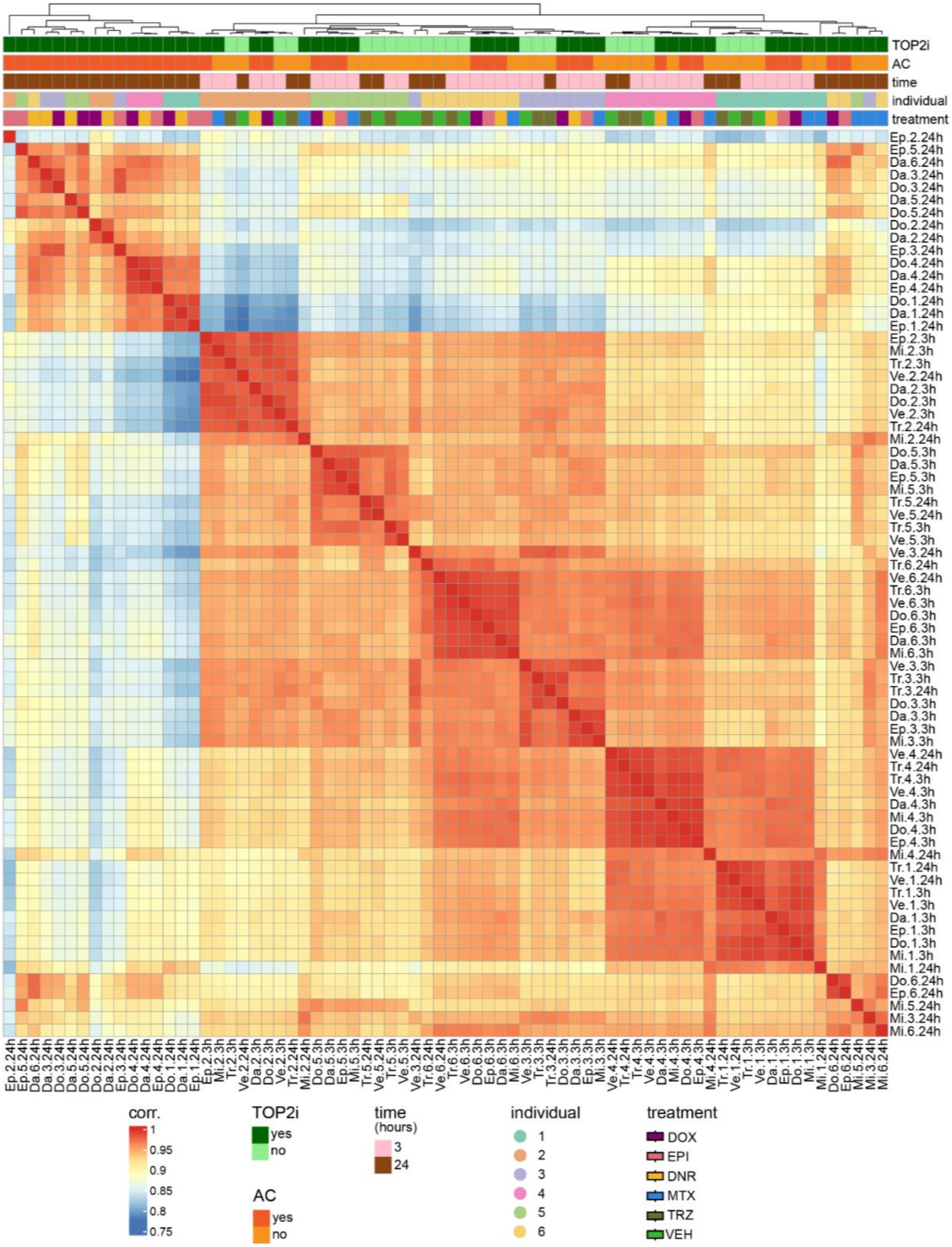


**Figure S6: RNA-seq samples cluster by treatment type, timepoint, and individual.** Pearson correlation of $\log_2$ cpm values across all pairs of samples.

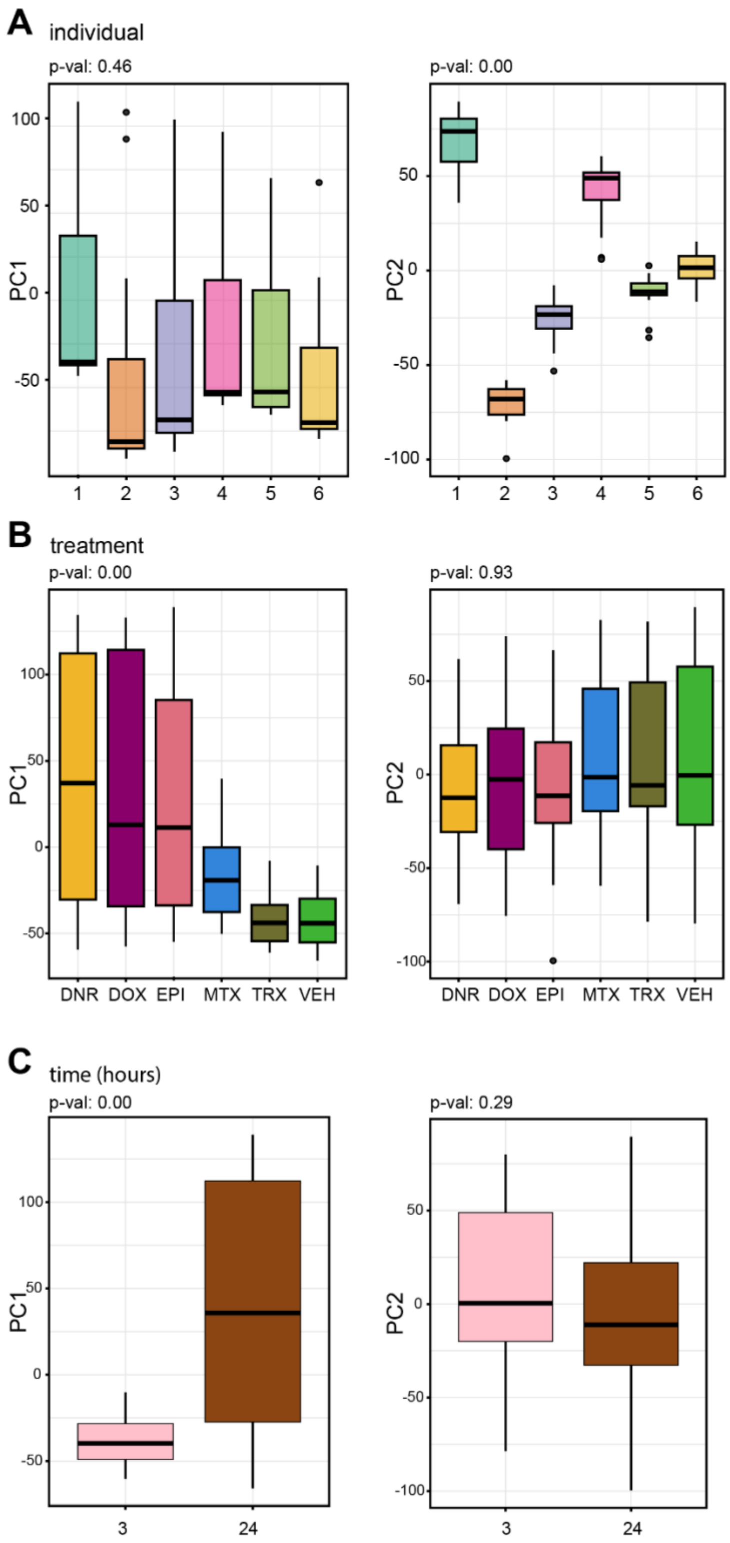


**Figure S7: PC1 associates with drug treatment and treatment time, while PC2 associates with individual.** Demonstration of variance contributed to the first two principle components from three major covariates in the study: individual, treatment, and time. **(A)** Variance of individual as a function of PC1 and PC2. The correlation between individual and each PC is calculated using a linear model. *P* values represent the significance of the F-statistic from the model. **(B)** Variance of treatment as a function of PC1 and PC2. **(C)** Variance of time as a function of PC1 and PC2.

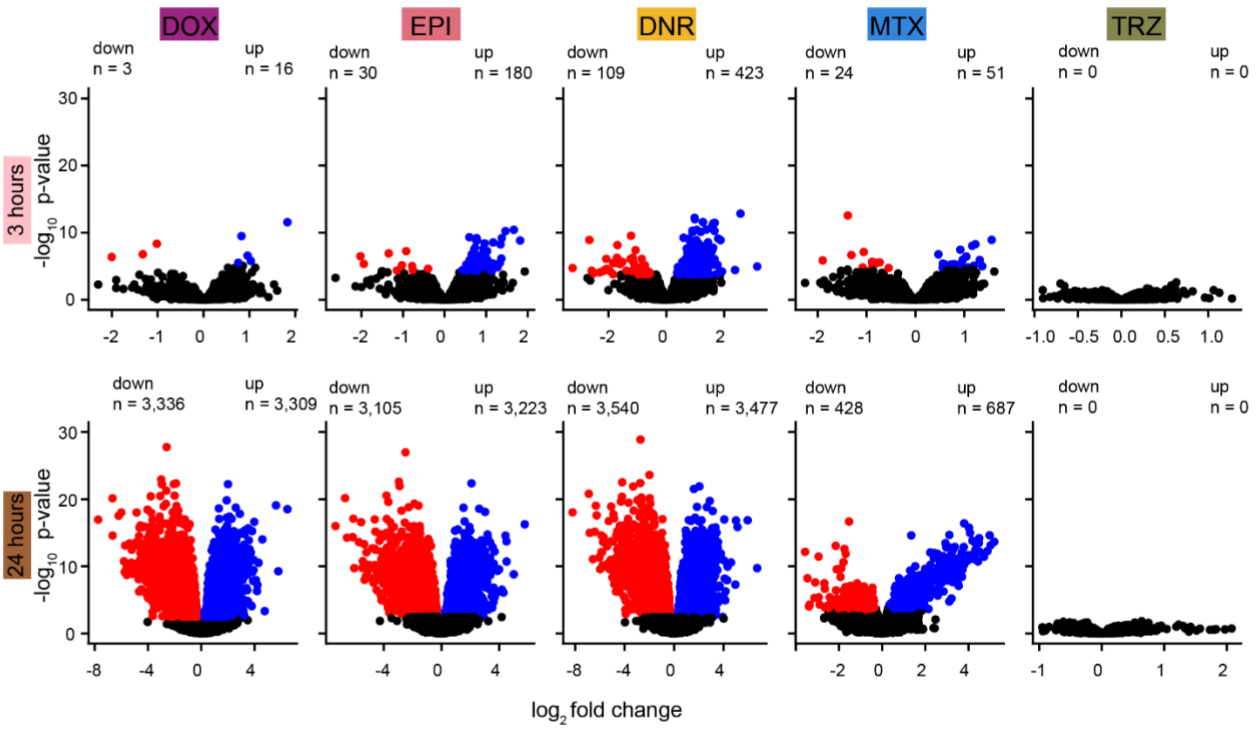


**Figure S8: Thousands of gene expression changes are induced in response to TOP2i treatment over 24 hours.** Volcano plots representing genes that are differentially expressed between drug and VEH treatment at each timepoint. Genes that are significantly up-regulated in response to treatment (adjusted *P* value < 0.05) are represented in blue, and genes that are significantly down-regulated are represented in red. The number of genes that are up- and down-regulated is given for each plot.

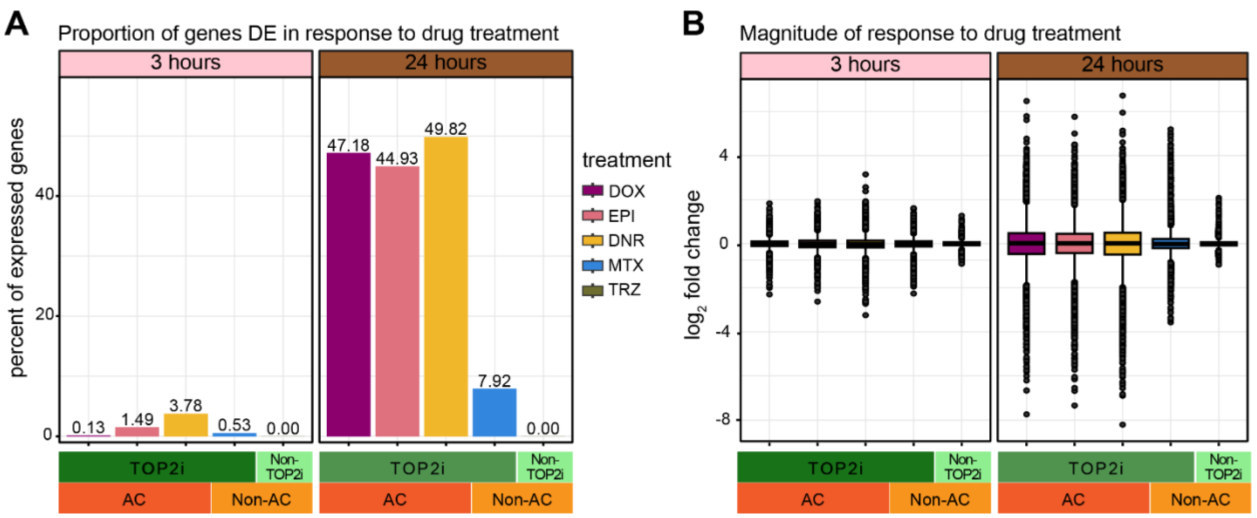


**Figure S9: ACs affect expression of nearly half of all expressed genes after 24 hours of treatment. (A)** Percentage of genes that are differentially expressed between each drug treatment and the VEH following three and 24 hours of treatment. **(B)** $Log_2$ fold change between drug-treated and VEH-treated samples for all 14,084 expressed genes following three and 24 hours of treatment.

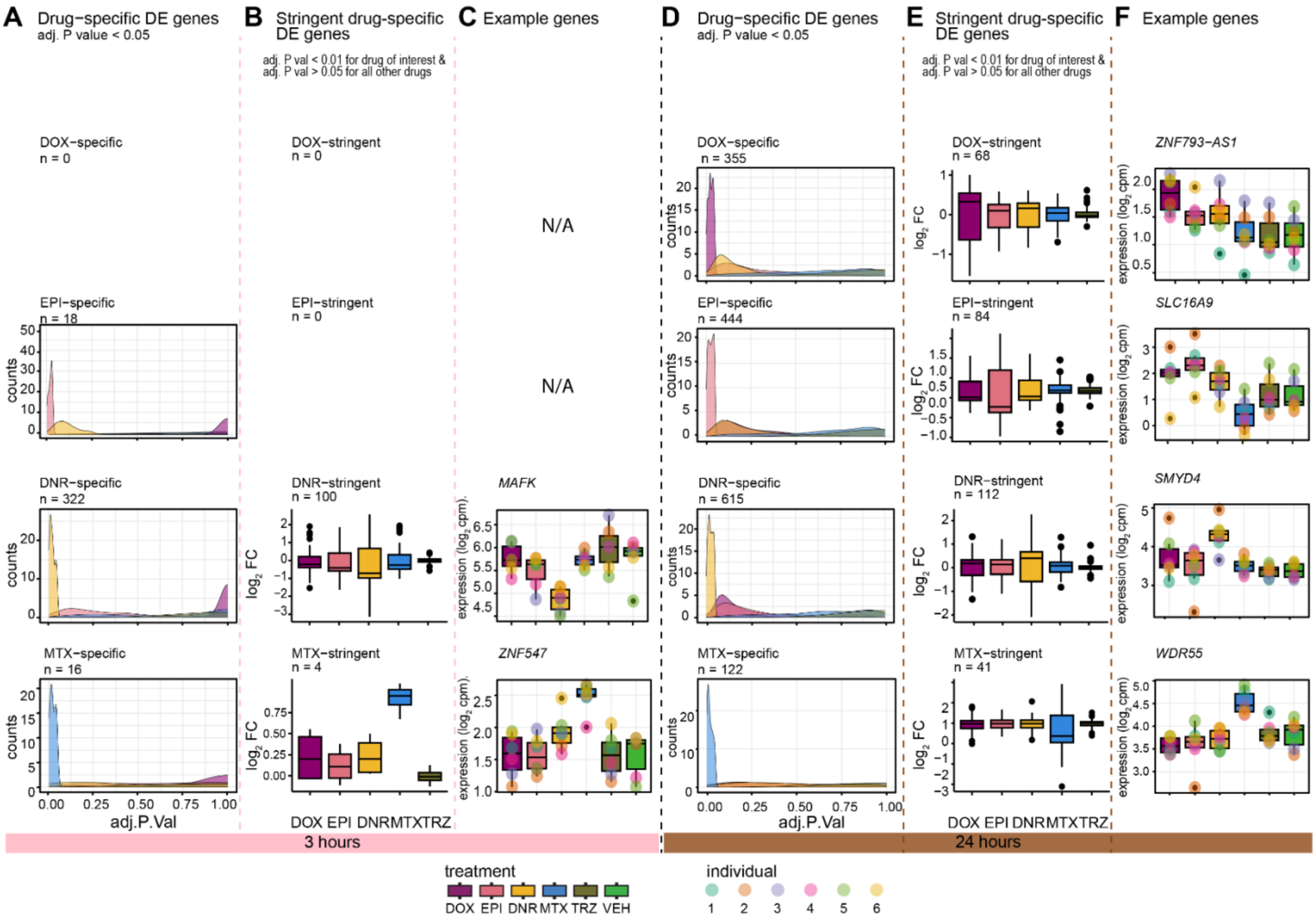


**Figure S10: A small number of genes respond to a single drug only. (A)** Distribution of adjusted *P* values for all genes classified as drug-specific response genes based on overlap of significantly differentially expressed genes meeting the adjusted *P* value cutoff of 0.05 across drugs at three hours. *P* values are shown for all drug treatments for the set of drug-specific genes (DOX: mauve; EPI: pink; DNR: yellow; MTX: blue; TRZ: dark green; VEH: light green). **(B)** The $\log_2$ fold change of all genes meeting a stringent adjusted *P* value cutoff of 0.01 for the drug of interest & > 0.05 for all other drugs to identify drug-specific response genes at three hours. **(C)** Examples of expression levels of stringently-identified drug-specific response genes across drug treatments including *MAFK* as a DNR-specific response gene and *ZNF547* as a MTX-specific response gene. **(D)** Distribution of adjusted *P* values for all genes classified as drug-specific response genes based on overlap of significantly differentially expressed genes meeting the adjusted *P* value cutoff of 0.05 at 24 hours. **(E)** The $\log_2$ fold change of all genes meeting a stringent adjusted *P* value cutoff of 0.01 for the drug of interest & > 0.05 for all other drugs to identify drug-specific response genes at 24 hours. **(F)** Examples of expression levels of stringently-identified drug-specific response genes across drug treatments including *ZNF793-AS1* as a DOX-specific response gene, *SLC16A9* as a EPI-specific response gene, *SMYD4* as a DNR-specific response gene, and *WDR55* as a MTX-specific response gene.

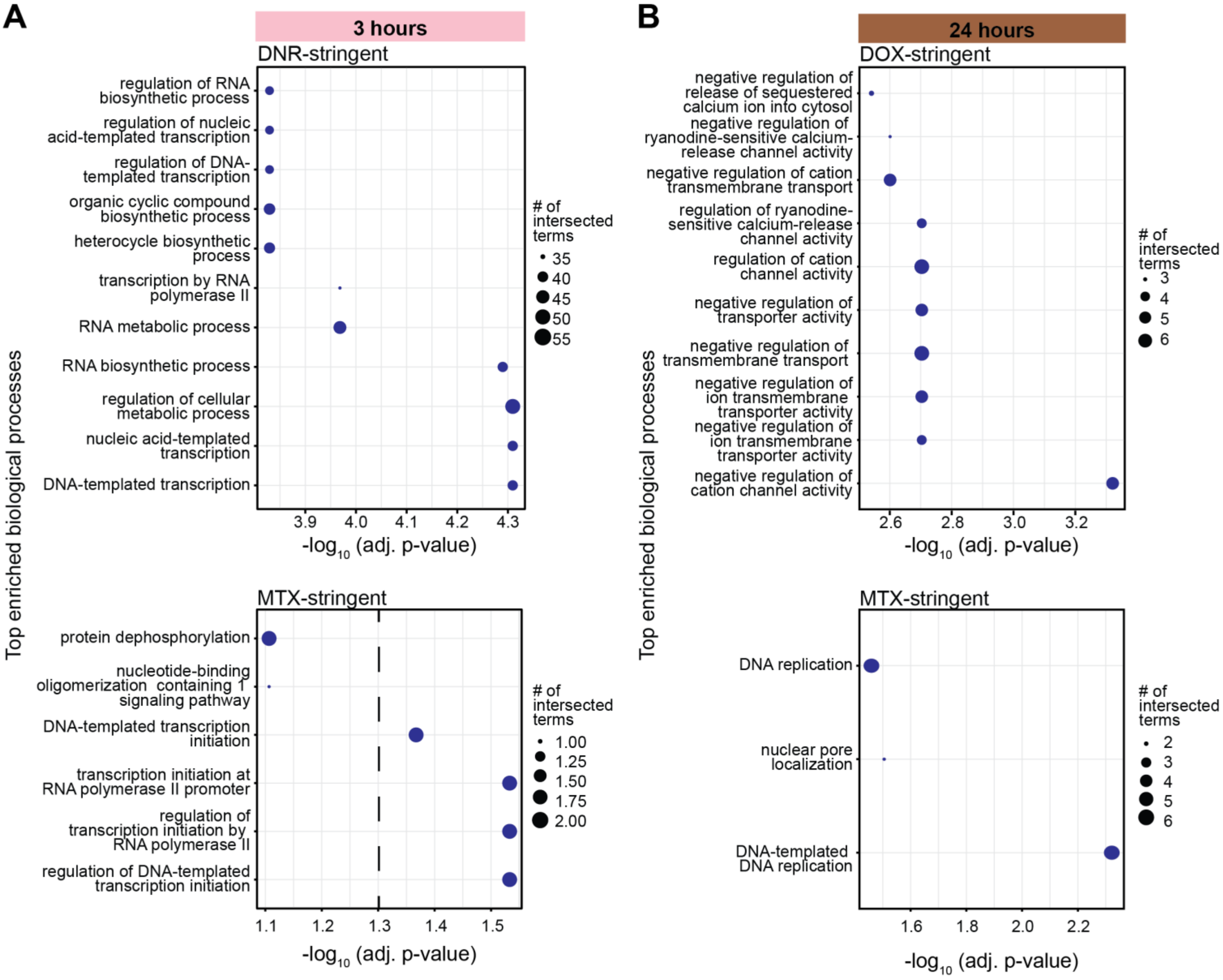


**Figure S11: Stringently-identified drug-specific response genes are enriched in biological processes. (A)** Biological processes enriched amongst genes classified as stringent DNR-specific and stringent MTX-specific response genes compared to all expressed genes following three hours of treatment. The top ten most enriched biological processes from Gene Ontology analysis that meet an adjusted *P* value cutoff of 0.05 are shown except for MTX-stringent where only processes to the right of the dashed line are significantly enriched. Dot size represents the number of stringent drug-specific response genes that are annotated as belonging to the particular biological process. There are no DOX-specific or EPI-specific response genes that pass the stringent threshold at three hours. **(B)** Biological processes enriched amongst genes classified as stringent DOX-specific and stringent MTX-specific response genes compared to all expressed genes following 24 hours of treatment. There are no EPI-specific or DNR-specific response genes that pass the stringent threshold at 24 hours.

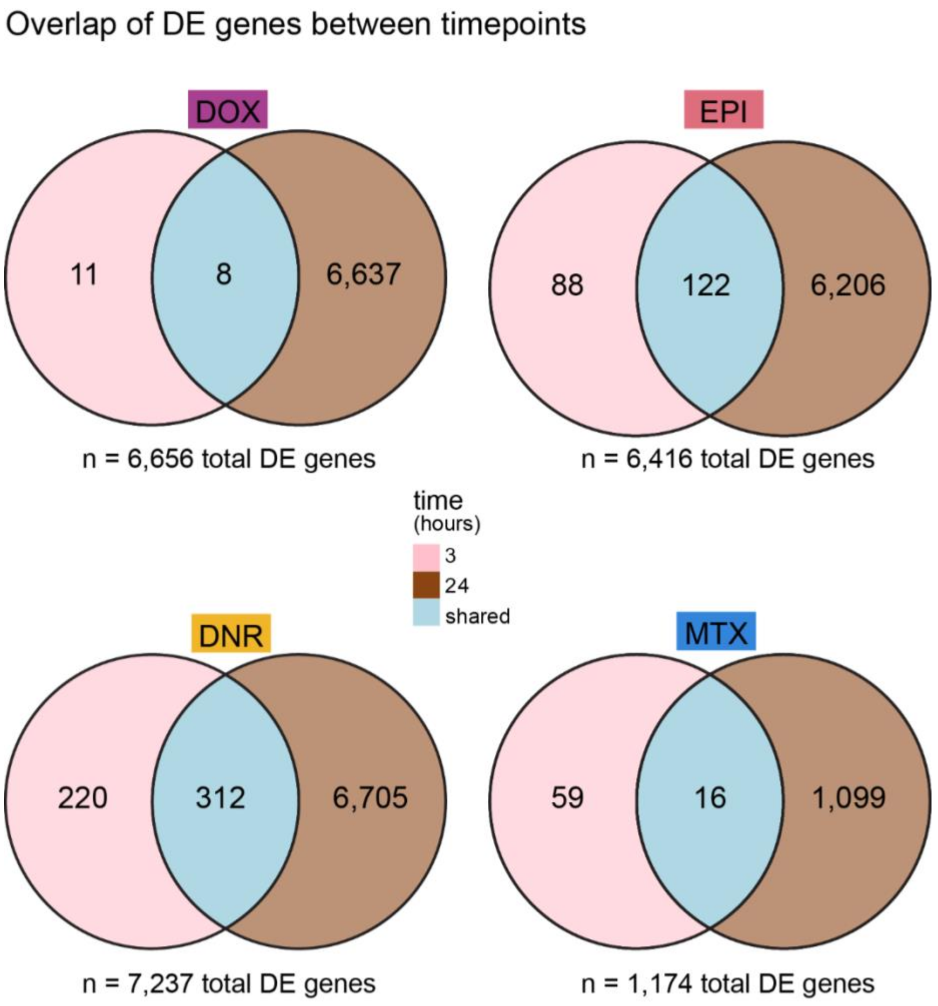


**Figure S12: Most genes that are differentially expressed in response to treatment at three hours are also differentially expressed at 24 hours.** Overlap of genes that are differentially expressed in response to each drug treatment at each time point. n = the total number of differentially expressed genes in response to the drug treatment across time points. The number of genes that are differentially expressed only following three hours of treatment is shown in pink, the number of genes differentially expressed only after 24 hours is shown in brown, and the number of genes differentially expressed at both timepoints is shown in blue.

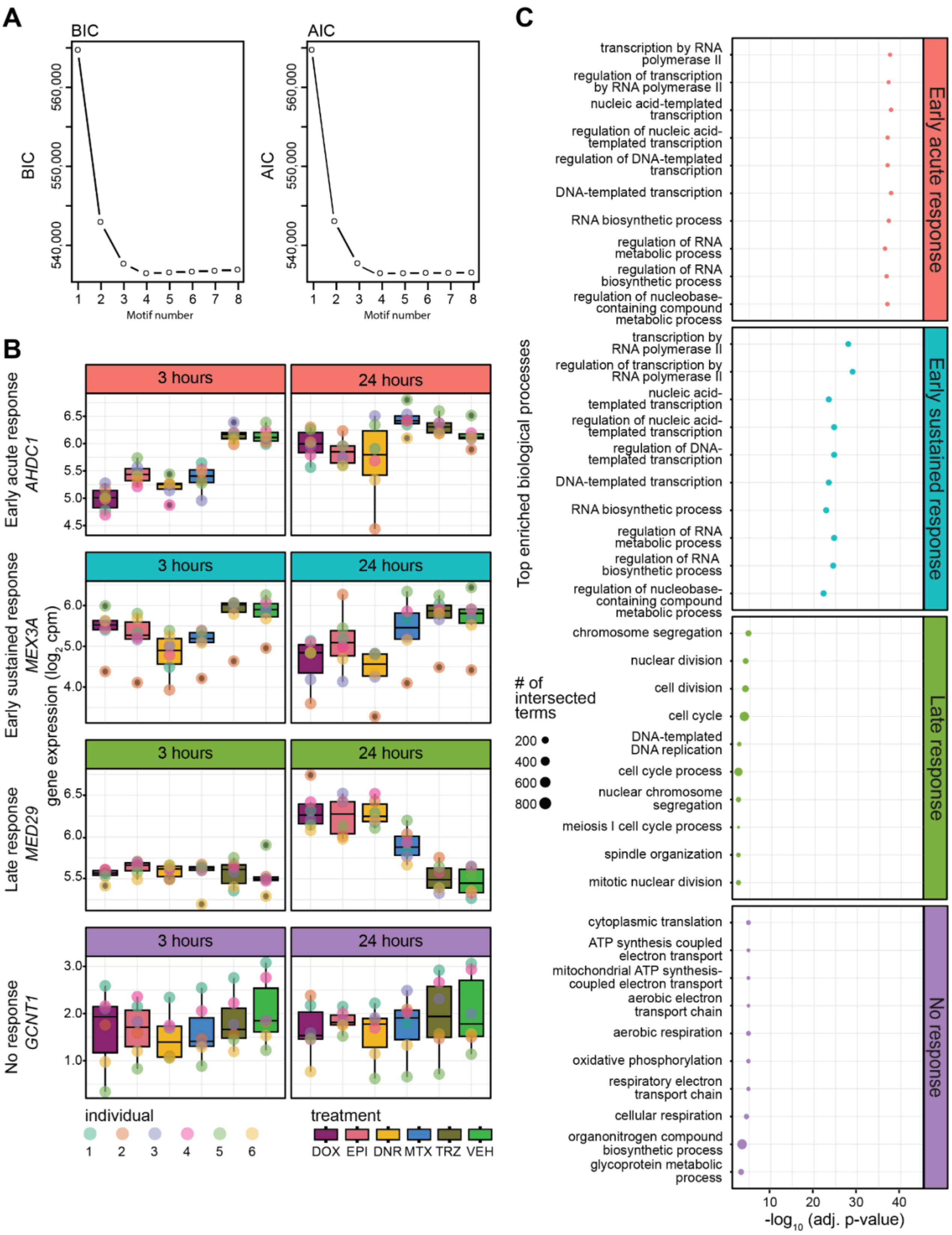


**Figure S13: Four gene expression signatures capture the response to TOP2i over time. (A)** Bayesian information criterion (BIC) and Akaike information criterion (AIC) at increasing numbers of Cormotif correlation motifs following joint modeling of pairs of tests. **(B)** Gene expression levels of genes assigned to each TOP2i response signature in each drug treatment at each time point. The *AHDC1* gene represents the Early-acute response motif (red), the *MEX3A* gene represents the Early-sustained response (blue), the *MED29* gene represents the Late response (green), and *GCNT1* represents the No response motif (purple). **(C)** The top ten most enriched biological processes (adjusted *P* value < 0.05) that are enriched in the response gene categories compared to all expressed genes. Dot size represents the number of correlation motif genes that are annotated as belonging to the particular biological process.

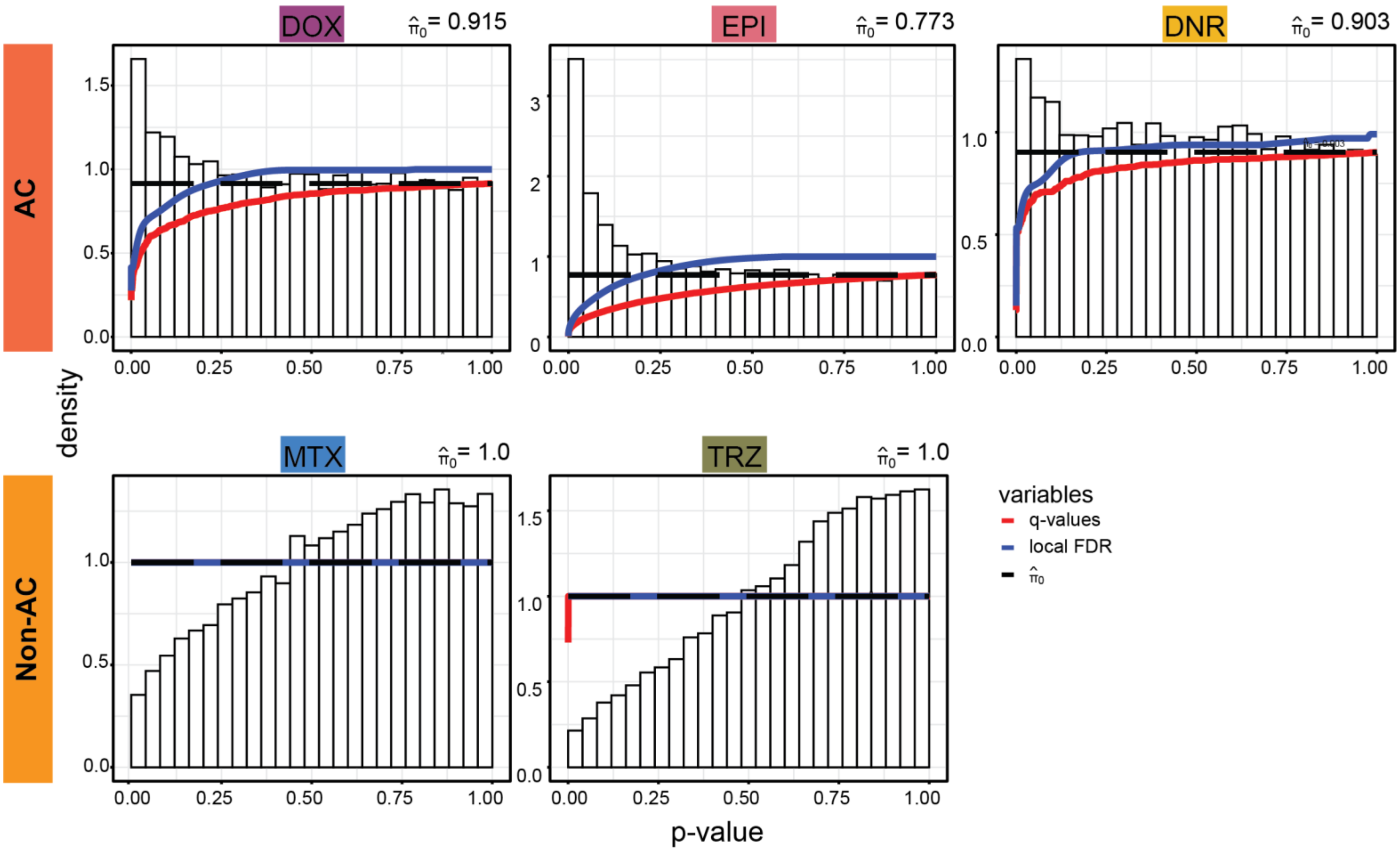


**Figure S14: AC treatments induce a small number of gene expression variation changes across individuals.** A density histogram of *P* values obtained from testing for differences in the variance between each drug treatment and the VEH control at 24 hours using an F-test. Q-values, local FDR and $\hat{\pi}_0$ (the estimated proportion of null tests in each distribution) are also shown.

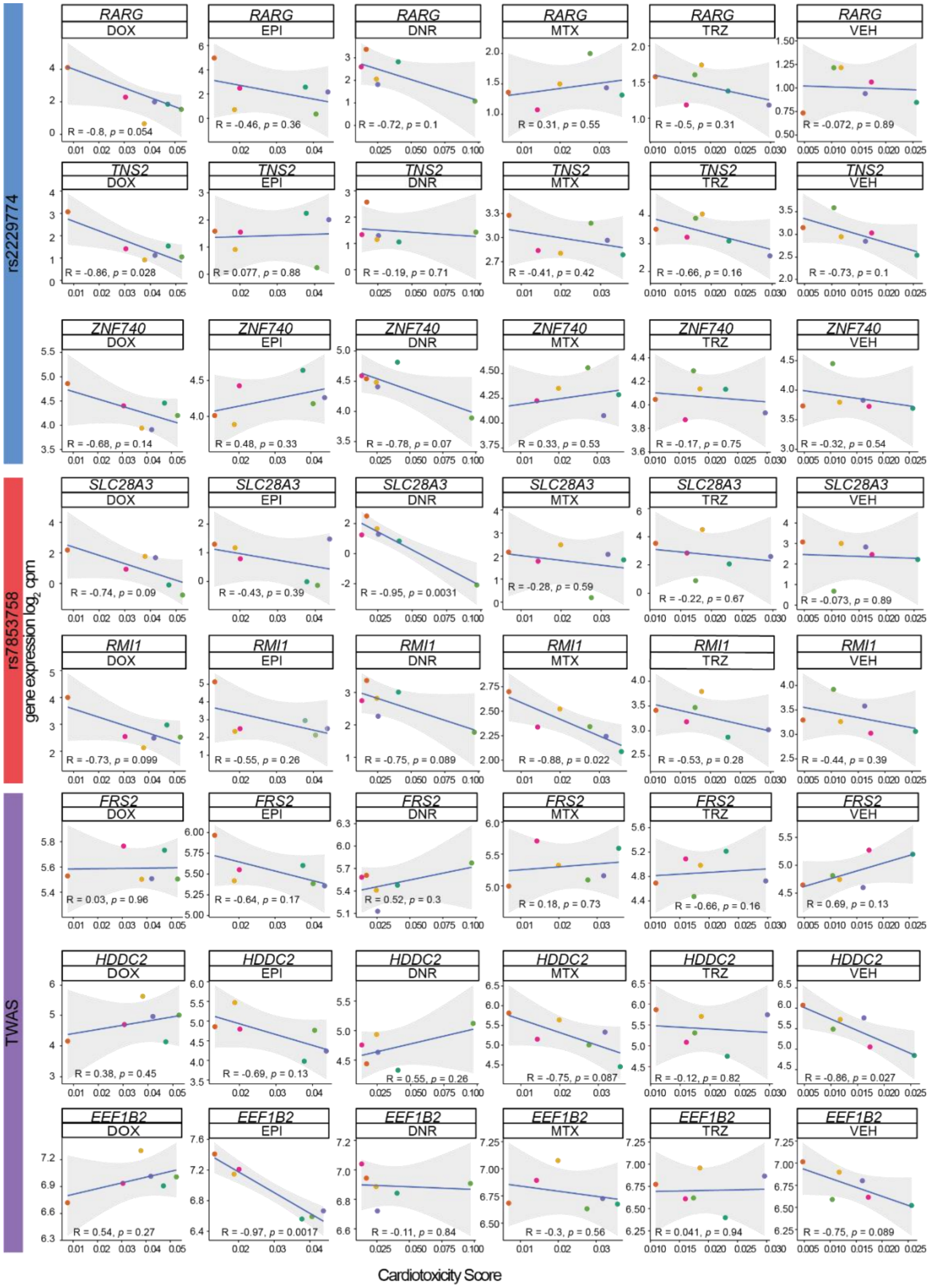


**Figure S15: TOP2i treatments induce expression changes in some genes that correlate with cardiotoxicity.** Correlation between gene expression levels of genes in AC-induced cardiotoxicity-associated loci, and a measure of cardiotoxicity across six individuals. A cardiotoxicity score was calculated by averaging the levels of lactate dehydrogenase and Troponin I. Pearson correlation values were calculated for each gene expression-cardiotoxicity pair across individuals.
